# What drives change? Characterizing scientific self-efficacy development in undergraduate research experiences

**DOI:** 10.1101/2025.08.18.670917

**Authors:** Qiyue Zhang, Paul R. Hernandez, Benjamin S. Listyg, Erin L. Dolan

## Abstract

Undergraduate research experiences (UREs) and course-based UREs (CUREs) promote students’ scientific self-efficacy growth. Yet, how self-efficacy develops during research is not understood. Furthermore, what students do during research varies in ways that likely affect self-efficacy development. We sought to address these knowledge gaps by collecting scientific self-efficacy data from CURE and URE students at nine universities at the beginning, middle, and end of a single term of research. We leveraged a theoretical advancement, latent state-trait theory-revised, to disaggregate the components of students’ self-efficacy into stable or trait-like self-efficacy and dynamic or state-like self-efficacy. We determined that students’ scientific self-efficacy was moderately stable during their research, with the most malleable component being beliefs in their abilities to figure out data collection and explain results. We also surveyed students ∼45 times throughout their research experience to test the extent to which research hours and types of research tasks contributed to self-efficacy development. We found that students who completed more analytic tasks experienced significantly more self-efficacy growth than students who completed other types of tasks, while time spent on research was not influential. Our results illustrate the importance of engaging students in analytic tasks during CUREs and UREs for fostering their self-efficacy development.

**Highlight for table of contents:** Using latent state-trait theory-revised, we found that students’ scientific self-efficacy was more stable than malleable over one research term. Beliefs about data collection and explaining results were most dynamic. Conducting more analytic tasks fostered self-efficacy, while the time spent and completion of other tasks had no effect.

## INTRODUCTION

Undergraduate education reform efforts in the life sciences have called for the involvement of all biology learners in research (American Association for the Advancement of Science, 2011). Undergraduate research experiences (UREs) have been championed for promoting student learning and development and for attracting and training the next generation of science researchers. Widespread enthusiasm for undergraduate research has been driven by evidence that research experiences afford distinctive opportunities for students to engage in the practices of science and learn to think and work like scientists (Gentile et al., 2017; Laursen et al., 2010; Lopatto & Tobias, 2010). The undergraduate life science community has responded to demands for more research opportunities by integrating research into undergraduate coursework, known as course-based UREs (CUREs). Research on both CUREs and UREs indicates that these educational environments support students pursuit of and persistence in science-related paths (Hanauer et al., 2017; Hernandez et al., 2018; Rodenbusch et al., 2016). Despite the widespread availability of UREs and CUREs in the life sciences, there is relatively limited understanding of how to design these experiences to maximize their effectiveness for students and only modest insight into the mechanisms through which research experiences foster desired outcomes for students (Auchincloss et al., 2014; Gentile et al., 2017; Linn et al., 2015).

Multiple theories have been leveraged to explore *how* research experiences shape students’ continuation in science, and the development of scientific self-efficacy beliefs (also called science self-efficacy and research self-efficacy) has emerged as an influential factor. Social cognitive career theory (SCCT; Lent et al., 1994) posits that an individual’s beliefs about their capabilities to perform career-related tasks (self-efficacy), coupled with their career interests and their beliefs about the consequences of their career choices (outcome expectations), drive their intentions to pursue particular careers (Lent & Brown, 1996, 2006). In the context of undergraduate research, SCCT predicts that engagement in research should foster students’ beliefs that they are capable of being successful in science, which in turn prompts them to set goals and engage in goal-directed behaviors consistent with continuing in science. Cross-sectional research informed by SCCT has shown that undergraduate researchers who report having greater research skills also express greater intentions to pursue graduate education and research-oriented careers, and this relationship was linked through gains in self-efficacy (Adedokun et al., 2013). Longitudinal research using SCCT has shown that greater levels of research experience predict higher identity as a scientist over time, and that this relationship is mediated by scientific self-efficacy (Robnett et al., 2015). While the short- and long-term associations between scientific self-efficacy and science-related outcomes appear clear, it is unclear how scientific self-efficacy develops during research experiences and what aspects of research experiences serve as sources of scientific self-efficacy.

### Longitudinal Development of Scientific Self-Efficacy during Research

Perhaps due to its role as a mediator in SCCT, little attention has been paid to how scientific self-efficacy develops over the course of a CURE or URE. Characterizing the longitudinal development of scientific self-efficacy within a CURE or URE is essential for understanding how to maximize the impact of these experiences. Longitudinal development can take many forms or patterns. Students’ scientific self-efficacy could grow steadily with experience, reflected in linear growth in average self-efficacy from timepoint to timepoint over a CURE/URE. Alternatively, students’ self-efficacy could grow early on in a CURE/URE, reflecting a steep “learning curve” evidenced by early non-linear gain and followed by a plateau. Finally, students’ self-efficacy could be relatively stable over a CURE/URE with context-dependent dips and spikes, which would appear as average self-efficacy that is unchanged over the CURE/URE, with fluctuations in the strength of a student’s self-efficacy from timepoint to timepoint. Research on undergraduate researchers’ scientific self-efficacy has primarily employed pre-post study designs and analytic methods that preclude distinguishing between these patterns (Frantz et al., 2017; Hess et al., 2023; Martin et al., 2021). Yet, characterizing patterns in students’ scientific self-efficacy over time is important because it would reveal whether self-efficacy gains are sustained over time or idiosyncratic to the timing of data collection (e.g., at the end of the experience following a culminating event like an end-of-term presentation). The resulting knowledge could also be used to inform the design of research experiences to maximize students’ self-efficacy growth.

### Theoretical Advancements that Enable the Study of Longitudinal Development

A recent theoretical development known as the latent state-trait theory-revised enables more accurate characterization of longitudinal development when coupled with a longitudinal study design (i.e., data collection at three or more timepoints). Latent state-trait-revised (LST-R) theory (Steyer et al., 2015). differs from classical test theory approaches because it provides a framework for analyzing the consistency, variability, and change of a latent variable over time. The use of LST-R as a framework involves fitting data to a series of alternative models of development, like the three hypothetical patterns of self-efficacy development described above, and determining the best-fitting model. Using LST-R to study the development of scientific self-efficacy is advantageous because a student’s rating of their scientific self-efficacy can be due to both static and dynamic factors. For instance, undergraduates’ scientific self-efficacy may be relatively stable, or “trait-like,” over the course of a research experience – either consistently higher if they have more prior experience or consistently lower if they have little prior experience. A student’s self-efficacy may also fluctuate based on the research task they are doing at a given moment (Jansen et al., 2020). If they are completing a familiar task that is relatively straightforward to accomplish, they may have momentarily higher self-efficacy. If they are completing a new task that is challenging to accomplish, they may have momentarily lower self-efficacy. Their self-efficacy may be further dampened if they fail at the task, they don’t have opportunities to try again or improve, or they hear messaging that they should be better at the task (e.g., an instructor or research mentor saying “other students have learned this task quickly – I don’t know why this has been a problem for you”). Disentangling how much of a student’s scientific self-efficacy rating (their “**state**” at a given point in time) is attributable to the student’s stable **trait** or to the dynamic “**occasion**” can advance our understanding of the dynamics of scientific self-efficacy, and offer insight into how day-to-day engagement in particular research tasks influences scientific self-efficacy development.

Historically, research on scientific self-efficacy in CUREs and UREs has been modeled using methods such as autoregression or confirmatory factor analysis (e.g., Adedokun et al., 2013; Hess et al., 2023; Lachance et al., 2020). Autoregressive models involve regressing a variable on its past value. These models offer the advantage of showing the degree to which prior levels of students’ self-efficacy influence future levels of self-efficacy (e.g., before and after a CURE/URE). However, autoregression does not account for measurement error, that is, the difference between their observed score on a latent variable (i.e., students’ average score on a self-efficacy scale) and their “true” score on that latent variable (i.e., their self-efficacy as a construct). Longitudinal confirmatory factor analysis (CFA) addresses this measurement issue by separating observed variability of scores into “true” variance and “error” variance at each timepoint. Yet, longitudinal CFA (and auto-regression for that matter) still conflate the occasion and trait elements of students’ scientific self-efficacy into a single “state” score, or their mean self-efficacy rating at a single timepoint. LST-R subsumes these traditional longitudinal models (Geiser, 2020) but moves beyond the traditional distinction between “true” and “error” variances in a construct by disaggregating a state score at each timepoint into an occasion component and trait component. This approach offers the added value of revealing which components of a latent variable are truly changing over time: the trait, which will endure over time, or just the occasion? LST-R makes this feasible by enabling the comparison of fit between different longitudinal measurement models to identify the optimal model that describes how developmental processes manifest. For example, if students’ self-efficacy is only due to a specific situation they experience and not on their stable traits, a state-only model will fit better than a model that incorporates traits and occasions. Alternatively, if scientific self-efficacy is more trait-like or contains a mixture of trait and occasion developmental processes, models that allow for traits and occasions will fit better than state-only models.

### Sources of Self-efficacy in Research Experiences

Many studies have shown that students in both CUREs and UREs increase in their scientific self-efficacy from pre to post experience (e.g., Dolan, 2016; Frantz et al., 2017; Hanauer et al., 2016, 2017; Hess et al., 2023; Martin et al., 2021; Newell & Ulrich, 2022; Wilczek et al., 2022). Yet, the specific aspects of research experiences that drive students’ scientific self-efficacy development are largely unknown. Bandura’s social cognitive theory offers an explanation for why students in CUREs and UREs experience scientific self-efficacy growth (Bandura, 1997). Social cognitive theory posits that individuals are agents with the ability to make choices, develop action plans, and engage in self-reflection through which they monitor their performance and reactions (Bandura, 2001). Efficacy, an individual’s belief that they can act on their choices and successfully execute their action plans, is core to individual agency (Bandura, 1977, 1997). Bandura (1997) theorized, and others have identified, four main sources of self-efficacy (Chen & Usher, 2013; Usher & Pajares, 2008). First, individuals rely on their prior experiences to make self-efficacy judgments. If they have been successful in their actions in the past, meaning they have had mastery experiences, they will judge themselves as efficacious. Second, individuals will assess their capabilities based on observations of the capabilities of others they perceive as like them, known as vicarious experience. Third, individuals will assess their capabilities based on feedback from “important others” (e.g., mentors, teachers, family, peers), which is known as social persuasion. Finally, an individual’s physiological or emotional state, or their positive or negative affect about an experience, will influence their efficacy.

Bandura asserted, and several studies have found, that mastery experience is the most influential source of self-efficacy (Bandura, 1997; Capa-Aydin et al., 2018; Chen & Usher, 2013; Usher & Pajares, 2008). A person’s sense of performance accomplishment is thought to be particularly influential because it is based on personal mastery experiences, meaning successful completion of a challenging task (Bandura, 1977). From an undergraduate research perspective, the training environment must afford opportunities to engage in research tasks, develop research skills, overcome obstacles, and eventually succeed. However, most research on CUREs and UREs treats them as a singular experience despite evidence that research experiences can vary widely (Ballen et al., 2018; Corwin et al., 2018; Gentile et al., 2017; Limeri et al., 2024).It is possible, even probable, that student gains vary as a function of prior experiences and mastery. For example, science students with limited experience conducting lab or field work may find all aspects of research challenging. They may realize gains in their scientific self-efficacy simply by spending time doing research, regardless of the specific tasks they carry out during research. Alternatively, science students with prior lab or field experience may find some tasks easier to accomplish (e.g., data collection) and others more challenging (e.g., posing an investigable research question). If so, they may experience scientific self-efficacy growth only when research experiences engage them in tasks that are new and more challenging to them, regardless of the amount of time they spend.

To our knowledge, no research has attempted to discern the research tasks that are necessary and sufficient for students to experience scientific self-efficacy growth. Rather, prior research on CUREs and UREs asked students to provide a global summary rating of their experiences, typically after the experience had concluded (Corwin et al., 2015; Gentile et al., 2017; Laursen et al., 2010). These methods are subject to the limitations of retrospective recall and autobiographical memory (Stone et al., 1999). For example, in gauging their experiences, students may favor recent events, such as presenting at a local symposium or national conference, which is often a positive culminating experience. Asking students to report about their research experiences only afterward misses the dynamic nature of day-to-day experiences (Shiffman et al., 2008) and may miss important features of research experiences that promote mastery. No studies of CUREs and UREs have connected students’ research tasks, that is, their day-to-day research activities, to changes in scientific self-efficacy. This raises questions about whether students must engage in specific types of research tasks, such as posing their own research question, or whether students must complete a certain number, range, or frequency of research tasks to build scientific self-efficacy.

### The Current Study

The current study aimed to address these knowledge gaps about students’ scientific self-efficacy development during research experiences by addressing the following research questions:

1. To what extent is scientific self-efficacy stable, or trait-like, versus situation-dependent in the context of undergraduate research?
2. Does the amount of time students spend on research tasks or the variations in the research time influence students’ situation-dependent scientific self-efficacy?
3. To what extent do different types of research tasks influence students’ situation-dependent scientific self-efficacy over and above time spent on research?

To accomplish this, we employed experience sampling, also known as ecological momentary assessment (EMA; Shiffman et al., 2008; Stone & Shiffman, 1994), to query CURE and URE students about their research tasks over one semester or summer at a diverse group of nine universities in the U.S. (see Maldonado Mendez et al., in review, for details). We texted survey links to query them ∼45 times over their research experience about the research tasks they engaged in that day, and the amount of time they spent on their research. We used correlated topic modeling to categorize their text responses into four main types of tasks: experimentation, analysis, communication, and instruction (Maldonado Mendez et al., in review). We also surveyed students at the beginning, middle, and end of their single term of research about their scientific self-efficacy.

To address our first research question, we compared a series of longitudinal models that distinguished “true” variability that is trait-like versus state-like. Our goal was to identify an LST-R longitudinal measurement model that optimally describes scientific self-efficacy development within a research experience. Then, we sought to determine the types of mastery experiences that influence scientific self-efficacy development. To accomplish this, we used research task types and time spent doing research as predictors of scientific self-efficacy development using the LST-R framework. We present the results of these analyses here.

## METHODS

The results reported here are part of a larger study of CUREs and UREs that was reviewed and determined to be exempt by the University of Georgia Institutional Review Board (PROJECT00000870).

### Participants

A total of 765 students (*N* = 765) completed data collection. The participants were nearly evenly split in terms of college rank; those in a CURE outnumbered those in a URE, and nearly half had no prior research experience. Most of the participants identified as women, almost one-third were from historically underrepresented groups, and a quarter were first-generation college students. Table 1 reports the complete details of the participants’ sociodemographics, institutional affiliations, research type, prior research experience, and year in school.

**Table 1.**
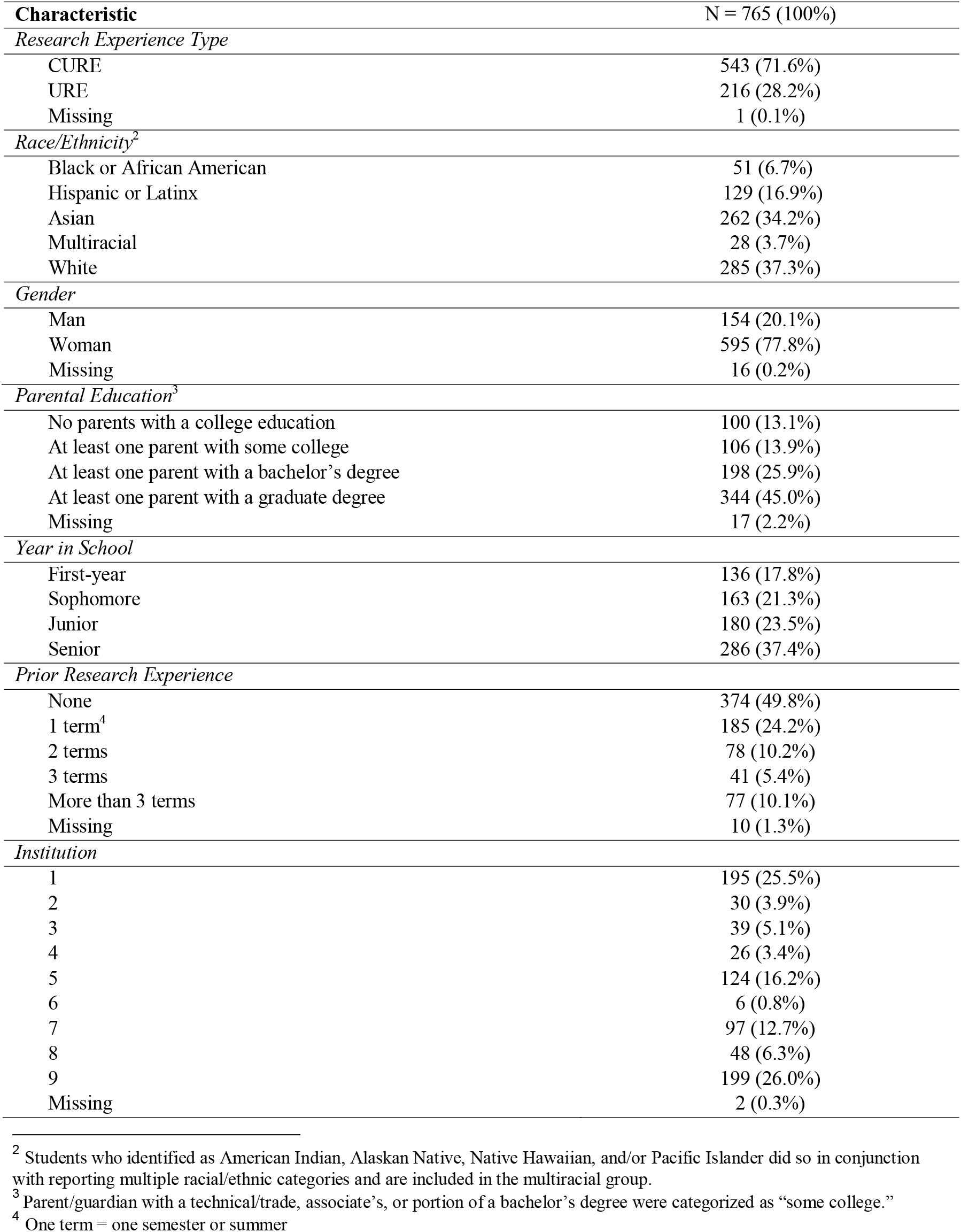
Student Characteristics.

Undergraduate students were recruited in the Spring, Summer, and Fall of 2021 from nine universities that varied in their research intensity and student populations (i.e., 6 doctoral public universities with very high research activity, 1 doctoral public university with high research activity, 2 public universities with Master’s level programs). Among the universities, one is classified as Historically Black, three as Hispanic serving, and two others as minority-serving. Recruitment methods are described in detail in (Maldonado Mendez et al., in review) and summarized here for context. The research team collaborated with liaisons at each university to recruit undergraduates enrolled in CUREs and UREs in the life sciences for the upcoming term using tailored panel management (Estrada et al., 2014). Students volunteered to participate in the study via a pre-survey linked to the study materials. Once enrolled, students were contacted by text message three times a week. Participants received survey links and compensation of $2 per response if their response rates stayed above 67%, resulting in ∼$100 compensation for the term.

### Data Collection

Data were collected using two main approaches: pre/mid/post-surveys distributed by email and EMA surveys distributed by text message. Methods and measures for each approach are described briefly below, and additional details are provided in the Supplemental Materials.

#### Pre/mid/post surveys

Recruitment surveys were distributed to potential participants in CUREs and UREs via institutional liaisons at our nine partner institutions. These surveys asked students to report on their scientific self-efficacy, terms of prior research experience (semester or summer), CURE/URE affiliation, course/section schedule, year in school, gender, race, ethnicity, and highest level of education completed by up to three parents/guardians. Follow-up surveys assessing scientific self-efficacy were sent at the midpoint and end of the research experience to examine change over time.

#### EMA surveys

Automated text messages were used to gather data about each student’s research experience as it unfolded. The Qualtrics^XM^ survey platform was used to send text messages with EMA survey links three times a week. Each survey was sent within 30 minutes after a student’s research experience was scheduled to conclude for the day (corrected for local time zone) and remained available for 30 minutes for that student to complete. This timeframe was established to ensure that students’ research experiences were fresh in their minds as they responded. The survey was structured with skip and display logic as follows: Students were queried about whether they did research that day. If they selected “no,” they exited the survey. If they selected “yes,” they were then asked a series of follow-up questions describing the amount of time and specific tasks they engaged in during their research that day. Additional questions related to students’ decision-making and collaboration were asked, although these are not the focus of the current study and thus are not included here. See Supplemental Materials for the full EMA survey.

### Measures

See Table 2 for the descriptive statistics for all measures.

**Table 2.** Descriptive statistics for outcomes and predictors at each timepoint.

|  | *n* | *M* | *SD* | *SK* | *KR* |
| --- | --- | --- | --- | --- | --- |
| Self-efficacy<sup>T1</sup> | 765 | 3.89 | .99 | -.36 | 2.98 |
| Self-efficacy<sup>T2</sup> | 693 | 4.22 | .91 | -.41 | 3.08 |
| Self-efficacy<sup>T3</sup> | 594 | 4.65 | .83 | -.64 | 3.75 |
| Research Hours<sup>T2</sup> | 718 | 1.35 | 1.12 | 2.32 | 10.44 |
| Research Hours<sup>T3</sup> | 756 | 1.17 | 1.09 | 2.42 | 10.45 |
| Research Hour Variation<sup>T2</sup> | 718 | 1.28 | .61 | .92 | 4.25 |
| Research Hour Variation<sup>T3</sup> | 754 | 1.27 | .66 | .89 | 4.55 |
| % Analysis<sup>T2</sup> | 715 | .24 | .25 | 1.04 | 3.56 |
| % Communication<sup>T2</sup> | 715 | .29 | .27 | .75 | 2.68 |
| % Experimentation<sup>T2</sup> | 715 | .28 | .32 | .97 | 2.68 |
| % Instruction<sup>T2</sup> | 715 | .19 | .21 | 1.23 | 4.33 |
| % Analysis<sup>T3</sup> | 736 | .21 | .24 | 1.21 | 3.98 |
| % Communication<sup>T3</sup> | 736 | .19 | .25 | 1.48 | 4.60 |
| % Experimentation<sup>T3</sup> | 736 | .27 | .33 | .98 | 2.58 |
| % Instruction<sup>T3</sup> | 736 | .33 | .28 | .53 | 2.34 |

**Notes:** SK = skewness, KR = kurtosis. % = proportion of each type of research task students reported.

#### Scientific Self-Efficacy (Outcome)

Scientific self-efficacy was measured using the scale developed by Estrada and colleagues (2011) to capture students’ confidence in their ability to carry out research-related tasks. Response options were on a 6-point scale from 1 (“Not confident at all”) to 6 (“Completely confident”). See Supplemental Materials S1 for the full scale.

#### Research Hours

Participants self-reported how many hours they had engaged in research that day (rounded to the closest half-hour). Participants responded to this question up to three times per week for the entirety of their CURE/URE. Responses were averaged over the first half of their research experience to operational research hours before the midpoint (*Research Hours*^*T2*^), and responses were averaged over the second half of their research experience to operational research hours before the end of the CURE/URE (*Research Hours*^*T3*^). Similarly, the standard deviation in the daily number of research hours over the first and second halves of the research experience operationalized variability in research hours (*Research Hours SD*^*T2*^ and *Research Hours SD*^*T3*^, respectively).

#### Types of Research Tasks

Participants who had engaged in research were asked to “Please describe what you did during your research today.” Their responses were recorded in a textbox with a 10-word minimum and could include multiple types of tasks. Participants responded to this question up to three times per week for the entirety of their CURE/URE. The daily types of research tasks were coded into four categories: Experimentation, Analysis, Communication, and Instruction (see Mendez et al., in review for details). Experimentation tasks involved preparing reagents and samples, carrying out techniques, and working with equipment. Analysis tasks involved a range of ways students worked with data, including collecting data, making measurements, calculating descriptive statistics, and using software. Communication tasks included formal and informal communication, including working on papers, posters, and presentations, and interactions with groups. Instruction tasks included attending meetings, lectures, or class, and completing training or assignments. Responses were summarized in terms of the percentage of each type of research task (to the total number of tasks) from the beginning to the midpoint (e.g., *%Analysis*^*T2*^) and from the midpoint until the end of the CURE/URE (e.g., *%Analysis*^*T3*^).

### Preliminary Analyses and Plan of Analysis

We conducted all analyses in Mplus 8.10 (Muthén & Muthén, 1998-2017). We ran a series of preliminary analyses to test the missingness mechanism, assess statistical assumptions, and diagnose outliers (see Supplemental Materials S2). Prior to addressing our research questions, we also assessed the descriptive statistics and bivariate correlations, which partially informed our expectations for the primary analyses (see Supplemental Materials S2). Next, prior to addressing substantive analyses, we conducted a preliminary assessment of the measurement equivalence of self-efficacy scores over time, which is a fundamental assumption of the longitudinal analyses required to address our research questions. The results indicated that longitudinal measurement equivalence was tenable (see Supplemental Materials S3 and Figure S1A-D for complete details).

To address Research Question 1 (i.e., *identifying the LST-R measurement model that provides optimal fit to scientific self-efficacy development*), we compared a series of six LST-R longitudinal measurement models that fit developmental processes that were primarily state-like, including (1) a State model, (2) a State Change model, and (3) an Autoregressive model (3), or that reflected mixture of states, traits, and occasions, namely (4) a Trait-State-Occasion (TSO) with a single trait, (5) a TSO model with multiple indicator-specific traits and an equal occasion variance model, and (6) a TSO model with multiple indicator-specific traits and unequal occasion variance. From this comparison, we identified a best-fitting model to address our other research questions based on Akaike Information Criterion and Bayesian Information Criterion (Akaike, 1974; Schwarz, 1978). See Supplemental Materials S5 and Figure S2 for a complete description of the alternative models.

To address Research Question 2 (i.e., *the influence of time spent or variations in time spent engaged in research tasks*), we predicted the state-like development of scientific self-efficacy from prior *Research Hours* and *Research Hours SD*. To address Research Question 3 (i.e., the *influence of different types of research tasks*), we predicted the state-like development of scientific self-efficacy from prior *%Analysis, %Classroom activities, %Communication*, and *%Experimentation*, controlling for *Research Hours* and *Research Hours SD*.

## RESULTS

### Students’ scientific self-efficacy is more stable than situation-dependent

We addressed our first research question (*To what extent is scientific self-efficacy stable, or trait-like, versus situation-dependent in the context of undergraduate research?*) in two ways. First, we took a descriptive approach to understanding the relative contributions of trait versus occasion-specific sources of self-efficacy by calculating a preliminary intra-class correlation coefficient (ICC). The ICC represents the proportion of “true” variance in self-efficacy that is trait-like, relative to the total “true” variance^1^. The ICC indicated 56% of the “true” variance in self-efficacy scores reflected stable trait-like differences among CURE/URE participants, while the remaining 44% occasion-specific. Next, we formally compared a set of six LST models to identify the model that best fit the development of self-efficacy over time in a CURE/URE (depicted in Supplemental Figure S2). We found that the trait-state-occasion (TSO) model with multiple indicator-specific traits (Figure 1) best fit the scientific self-efficacy data over time based on AIC and BIC indices (see Supplemental Table S4 for a comparison of the models). This result means that true state variability in students’ scientific self-efficacy is best modeled as a combination of a latent trait (stable over time) and latent occasions (fluctuates due to situational changes).

**Figure 1.**
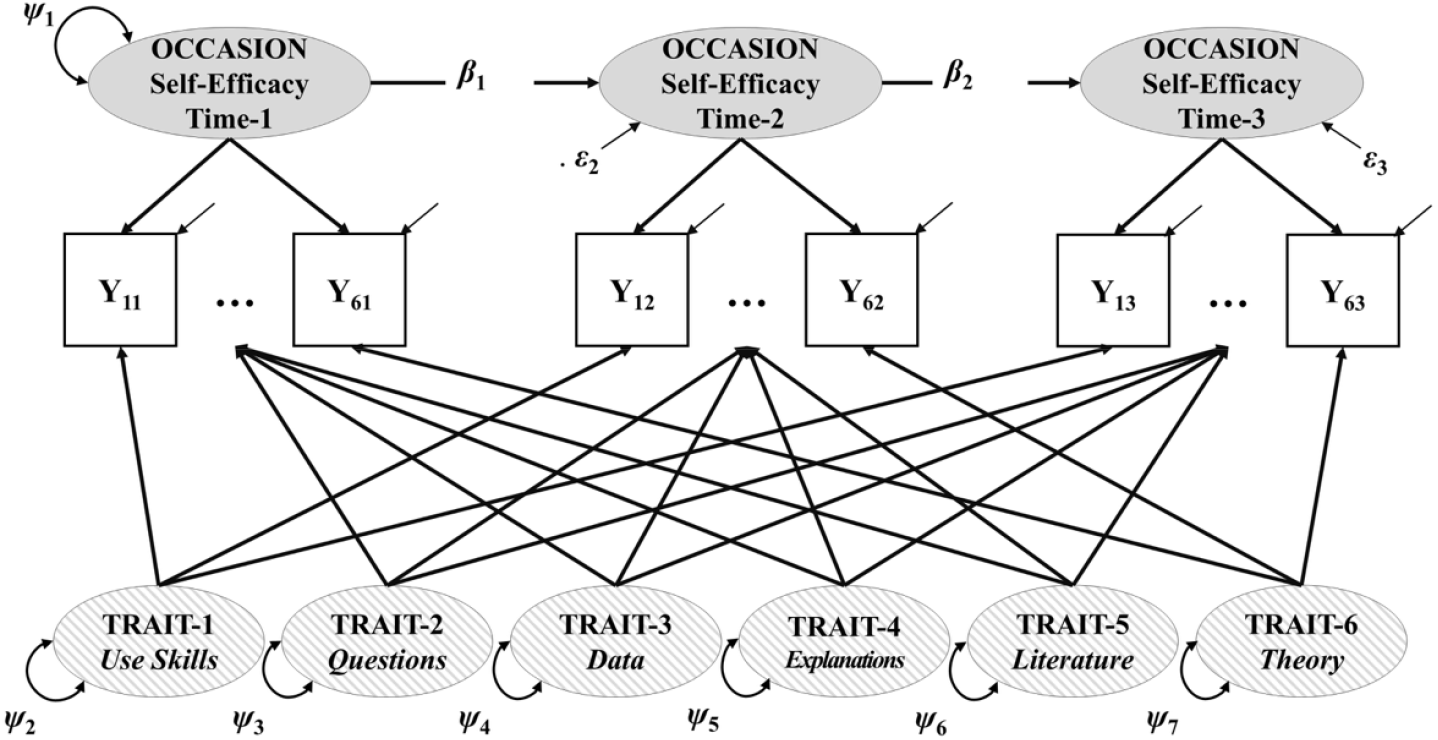
The Trait-State-Occasion (TSO) Model. The TSO model with multiple indicator-specific traits was the best-fitting among the six longitudinal models tested. This model disaggregates latent state into trait variability that is consistent across timepoints (patterned 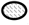), and occasion variability that fluctuates across timepoints (shaded 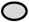). ψ = variances of latent trait or occasion variables. ε = residual variance of latent trait or occasion variables. β = autoregressive coefficients reflecting carry-over effects between occasions. Y_it_ = observed score for indicator *i* at time *t. e*_it_ = measurement error for indicator *i* at time *t*.

### Students’ beliefs about their data collection and explanation skills are most dynamic

Although the ICC provides valuable information about the sources of true variability in self-efficacy in aggregate (i.e., overall self-efficacy), it does not provide insight into which indicators of self-efficacy might provide levers for change or intervention. Our LST-R model enables indicator-level, or item-by-item, analysis to gain insight into which indicators are more situationally dependent and thus may provide opportunities for change through CURE/URE activities. Our indicator-level analysis revealed distinct patterns in stability over time (Supplemental Table S6). For example, we found that true variance in the “use technical science skills” indicator (Item 1) was the most stable and least dependent on the situation or timepoint (71% trait, 29% occasion-specific). This means that students’ beliefs about their ability to “use technical science skills” were least changed by their research experience over time, and that any changes in students’ global scientific efficacy were least influenced by their confidence to use technical science skills. In contrast, we found that true variance in the indicators “figure out what data/observations to collect and how to collect them” (Item 3) and “create explanations for the results of a study” (Item 4) were least stable (42% and 41% trait, respectively) and most dependent on the situation or timepoint (58% and 59% occasion-specific, respectively). This means that students were most dynamic in their beliefs about these abilities, and shifts on these indicators over time had the most influence on their global scientific self-efficacy development. Finally, we found that true variances in the “generate a research question to answer” (Item 2), “use scientific literature and/or reports to guide research” (Item 5), and “develop theories (integrate and coordinate results from multiple studies)” (Item 6) indicators were equally partitioned between trait and occasion (53%, 53%, and 52% trait and 47%, 47%, and 48% occasion-specific, respectively). This means that students’ beliefs about these abilities were close to equally composed of a stable component not influenced by their research experience and a dynamic component influenced by the situation or timepoint.

### Engaging in Analysis tasks promotes situation-dependent scientific self-efficacy development

Having identified the best-fitting longitudinal measurement model (i.e., TSO; Figure 1), we aimed to address our second question: *Does the average time students spent on research or variability in the time they spent affect their occasion-specific scientific self-efficacy?* We found that neither the average time students spent on research nor the variability in the time they spent affected their occasion-specific level of self-efficacy. Closer examination of the correlation and regression results (see Supplemental Materials) revealed that standard deviation in time spent on research is consistently marginal in positively predicting scientific self-efficacy at mid-term. This result may suggest that variation in time spent on research may be advantageous for students’ self-efficacy growth.

We also used our TSO model to address our third research question: *Do different types of research tasks influence students’ situation-dependent scientific self-efficacy over and above time spent on research*? To accomplish this, we assessed the influence of each type of research task (i.e., Experimentation, Analysis, Communication, Instruction) while controlling for prior scientific self-efficacy, average time spent on research, and variability in time spent on research in all models. We modeled each task type separately to avoid suppression effects (see Supplemental Materials). We found that students who engaged in a greater percentage of *Analysis* tasks reported slightly higher scientific self-efficacy at both the midpoint and the end of their CURE/URE (Figure 2; β = 0.12 and 0.13, respectively). We also found that, in assessing potential effects of Experimentation tasks, higher *variability in the time spent on research* predicted slightly higher scientific self-efficacy at the midpoint. However, this effect is not salient when the outliers are removed from the model (Supplemental Table S7).

**Figure 2.**
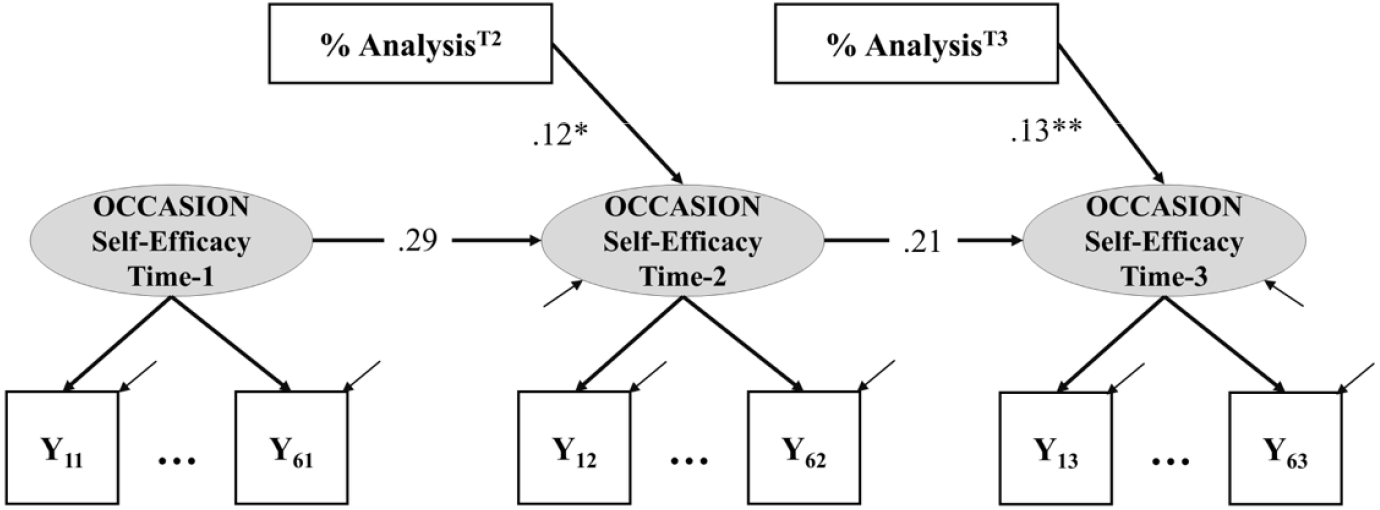
Engaging in Analysis tasks fosters students’ scientific self-efficacy development. Engaging in Analysis tasks significantly predicts students’ occasion-specific scientific self-efficacy over time, controlling for total hours and variability in research time. This effect is depicted here as a structural model with statistically significant structure coefficients between Analysis tasks and occasion self-efficacy timepoints 2 and 3. * *p*<.05, ** *p*<.01.

## DISCUSSION

Most prior research on undergraduate research, including both CUREs and UREs, measures students’ scientific self-efficacy at the start and end of an experience. These pre-/post-designs provide only limited information about developmental processes, such as how students’ self-efficacy changes over time and what factors may influence these changes. In this study, we collected data on students’ scientific self-efficacy at three timepoints and leveraged the LST-R framework to characterize how students’ scientific self-efficacy changes within a semester-long research experience. Importantly, we found that the trait-state-occasion (TSO) model fit the data best, indicating that undergraduate researchers’ scientific self-efficacy development is both trait-like and situation-dependent. Furthermore, we observed that students’ self-efficacy is moderately stable over a semester of research; only ∼44% of the true variance in their self-efficacy is dynamic. This pattern could only be identified using an LST-R type model of longitudinal data. The stable, trait-like component suggests that students are coming to research experiences – both CUREs and UREs – with prior experiences that have shaped their scientific self-efficacy in ways that are unaffected by a single semester of research experience. These students likely come to research with personal experiences that have fostered high scientific self-concept (e.g., being raised in a family with scientists or one that placed value on science experiences) or with prior academic experiences through which they learned to use science skills (e.g., lab instruction) (Jansen et al., 2015).

Despite the relative stability of students’ self-efficacy, our LST-R model made it possible to identify self-efficacy beliefs that were more malleable and those that were more crystallized at this stage of development. For instance, we found that “using technical skills” was the most stable among the self-efficacy beliefs of students in our study. This result is somewhat surprising given that about half of the students in our study had no prior research experience, and most had less than one term of research experience. Thus, they are unlikely to have experience with the specific technical skills they are using in their CUREs/UREs. However, they likely had prior “demonstration” lab instruction in which they followed established protocols designed to complement didactic coursework and reinforce their understanding of well-established concepts. Completing demonstration lab exercises likely enabled students to develop technical skills along with confidence in their abilities to be successful in technical aspects of scientific work. In contrast, students’ beliefs about their abilities to figure out data collection and explain their results were most dynamic, presenting fruitful targets for intervention to boost student self-efficacy. The percentage of analysis tasks that students carried out was also the most consistent contributor to their self-efficacy development, as analytic tasks require students to make decisions about their data and interpret their results. In other words, students who reported more analytic tasks also experienced greater self-efficacy growth, specifically because they grew in their beliefs about their abilities to figure out what data to collect and how to collect it, and to create explanations for the results of a study. These results also suggest that students experience analytic tasks as a greater sign of mastery than experimentation, communication, and instruction tasks, perhaps because they perceive analytic tasks to be the most difficult (Honicke et al., 2023).

This result raises questions about why students differed in their engagement in analytic tasks in the first place. Analytic tasks included working with data, graphs, statistics, and software, as well as scoring and making measurements (Maldonado Mendez et al., in review), all of which are standard in many research experiences in the life sciences. One possibility is that students who reported conducting more analysis tasks may have collected the data they needed and could shift from conducting techniques and completing protocols (experimentation tasks) to working with and making inferences from their data. Students may have experienced this shift as a sign that they had successfully performed the research and accomplished something scientifically (mastery experience), which fostered their self-efficacy. Students who experienced less research progress may have continued experimentation or engaged in communication or instruction without analysis. As a result, these students may have missed engaging fully in the scientific process, which limited their sense of mastery. This interpretation is consistent with the notion that experience (i.e., time spent on research) is not equivalent to mastery (Hirshfield & Chachra, 2019), which is further supported by our results showing that research time and research time variation were not predictive of students’ self-efficacy development over time.

It is important to note that the design or implementation of the CUREs and UREs may have limited students’ engagement in analytic tasks. Some research experiences primarily involve students in conducting mundane or menial tasks, such as repeatedly collecting the same data for extended periods (Limeri et al., 2019, 2024). Although research often demands this type of work, our results point to the importance of ensuring students have opportunities to move beyond experimentation tasks to more cognitively engaging analytic work. Furthermore, we cannot rule out alternative explanations of how analytic tasks foster students’ self-efficacy or alternative sources of self-efficacy in general. For instance, it may be that students whom their instructors or mentors viewed as more capable were given more analytic tasks to complete. In this situation, analytic tasks may function as social persuasion – a sign that their instructor or mentor thinks they are up to the task. It may also be that students who experienced more experimental failure and thus had to keep repeating experiments (i.e., reporting more experimentation tasks and fewer analytic tasks) lost confidence in their scientific abilities. Students who face more experimental failure may experience more negative emotions, such as stress associated with things going wrong or disappointment with things not working, which dampens their self-efficacy (Corwin et al., 2022). It may be that it is easier to “succeed” at analytic tasks because analysis will generate a product regardless of its quality, while experiments that fail may not generate any product. Future research could test these alternative mechanisms by measuring students’ research tasks and self-efficacy, along with other sources of self-efficacy, over time.

Collectively, our results have implications for the design and implementation of CUREs and UREs to maximize students’ self-efficacy growth. First, the time spent on research is less important than what students do during this time. This means that CUREs and UREs that involve students in a few hours of research each week over a semester may be sufficient to foster self-efficacy growth. More hours may not be necessary or beneficial, especially if students have competing demands on their time because of academic and personal commitments. This recommendation is consistent with evidence-based learning strategies, including spacing practice (Donovan & Radosevich, 1999; Rohrer, 2015) by doing research a few hours per week for many weeks and “interleaving” (Rohrer, 2012) by mixing up research tasks rather than focusing on a single type of task (e.g., experimentation). Second, research instructors and mentors can bolster self-efficacy by ensuring students have ample opportunities to make decisions about data collection and make sense of results. Notably, students in our study, including those who were new to research, reported completing analytic tasks in both the first and second halves of the semester. Instructors and mentors should strategically design research experiences to engage students in analytic tasks throughout their research experience, such as by identifying how to make observations or collect data before doing so themselves and explaining how published results relate to or inform the research they are doing (see Mendez et al., in review, for more examples).

## LIMITATIONS

Several aspects of our study limit what can be concluded. First, we assigned a single task type to each student’s description of their research on a given day to measure their research tasks. It may be that students completed multiple types of tasks on a single day and devoted different amounts of time to each task. Future research should explore the affordances and constraints of more fine-grained measurement of research tasks (e.g., all the types of tasks completed on a given day, how much time was spent on each type of task). Second, we opted to focus our study of scientific self-efficacy on tasks as an operationalization of mastery experience as a source of self-efficacy. To more fully test the potential for self-efficacy theory to explain students’ development of scientific self-efficacy through research experiences, other sources of self-efficacy (i.e., social persuasion, vicarious experience, emotional arousal) should be measured alongside the completion of research tasks. Third, the students in our study were all enrolled at universities and participated in the study for one semester. Thus, our results may not be reflective of the students’ self-efficacy development in other institutional settings (e.g., community colleges) or across multi-semester research experiences. Future research on students’ self-efficacy development during CUREs and UREs in these and other contexts should leverage study designs and modeling methods that allow for developmental insights such as those gleaned from our research (e.g., data collection over three or more timepoints, modeling that disaggregates state and trait variability).

## ACKNOWLEDGMENTS

We are grateful to our participants for sharing their time and experiences and to the site coordinators at each of the nine universities in the study. This work was supported in part by National Science Foundation EHR Core Research Grant 1920407 and the Georgia Athletic Association Professorship for Innovative Science Education. Any opinions, findings, conclusions, or recommendations expressed in this material are those of the authors and do not necessarily reflect the views of any of the funding organizations.

## Footnotes

1 ^1^ 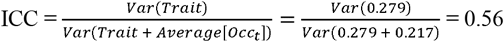 Where *Var* = Variance of a latent variable, *Occ*_*t*_ = latent self-efficacy on the occasion variable at time *t*, and *Trait* = latent trait self-efficacy across all timepoints. Variances were estimated in a preliminary simplified LST-R model as shown in Figure S1c, but without auto-regression paths.

