## Supplemental Materials for "What drives change? Characterizing scientific self-efficacy development in undergraduate research experiences"

Qiyue Zhang, Paul R. Hernandez, Benjamin S. Listyg, Erin L. Dolan

This supplement contains the following:


| **Section and Item** | **Page** |
|---|---|
| S1. Measures | 2 |
| S2. Preliminary Data Analyses | 3 |
| S3. Measurement Equivalence/Invariance | 4 |
| Table S1. Confirmatory factor analysis for self-efficacy at the individual timepoints | 4 |
| Table S2. Summary of model fit and measurement invariance test across CURE and URE groups at the individual timepoints | 5 |
| Table S3. Model comparison fit statistics for longitudinal invariance T1-T2-T3 | 5 |
| S4. Longitudinal Measurement Models (Figures S1A-D) | 6-8 |
| S5. Model Comparison (Figure S2) | 9-10 |
| Table S4. Measurement model comparison | 11 |
| S6. Results | 12 |
| Table S5. Correlation matrix of outcomes and predictors at each timepoint | 13 |
| Table S6. Consistency, occasion specificity, and reliability estimates for indicators (items) across time | 14 |
| Table S7. Summary of estimates from the final model | 15-16 |
| References | 16-17 |

**S1. Measures**

Scientific Self-Efficacy (Adapted from Chemers et al., 2011; Estrada et al., 2011)

Please rate your confidence on the ability to perform the following tasks on a scale from 1 (not confident at all) – 6 (completely confident).

Item 1: Use technical science skills (use of tools, instruments, and/or techniques).

Item 2: Generate a research question to answer.

Item 3: Figure out what data/observations to collect and how to collect them.

Item 4: Create explanations for the results of a study.

Item 5: Use scientific literature and/or reports to guide research.

Item 6: Develop theories (integrate and coordinate results from multiple studies).

Experience Sampling Measures

Did you do [research] today? – Yes/No

- If yes: How many hours? (to closest half hour)
- If yes: Please describe what you did during your research today (10 word minimum)
- If no, then asked: Are any of these reasons you did NOT do research today? Response options were multiple choice:
    - I was not scheduled to do research today.
    - I was too busy to do research today.
    - There were no research tasks for me to accomplish today.
    - I will do research later today.
    - Other (please explain)

**S2. Preliminary Data Analyses**

We conducted preliminary analyses to investigate the missingness mechanism and test the validity of statistical assumptions. We examined the missingness mechanism using Little's test (Little, 1988) and detected outliers using Mahalanobis distance. Interclass correlations (ICCs) and design effect values (DEFFs) were used to evaluate nesting effects. We assessed univariate normality using skewness, kurtosis, Shapiro–Wilk test and evaluated multivariate normality using the Doornik-Hansen test. In addition, we examined the relative homogeneity of variances and extreme collinearity of the outcomes using regression diagnostics (Kline, 2023).

The missing rate is 9.41%-9.67% for indicators at T2 and 22.35%-22.61% for indicators at T3. MCAR test indicated that a missing-complete-at-random assumption is plausible, $\chi^2(144)=153.05$, $p=.29$. We detected 25 outliers (3.27%) and excluded them from the final conditional model. In addition, we have found significant nesting effects by universities (ICCs=.02-.05, DEFFs=2.75-5.37). Thus, we included universities as dummy coded predictors in the final conditional model to control for the nesting effect. The univariate and multivariate normality were violated, $\chi^2(6)=95.81$, $p=<.001$. The homogeneity of variances was also violated based on the regression diagnostics. Thus, our analyses adopted the maximum likelihood estimation with a robust estimator. The variance inflation factors (VIFs) indicated no extreme collinearity in the predictors (VIFs=1.43-1.60).

### S3. Measurement Equivalence/Invariance

We investigated the internal structure and invariance properties of the self-efficacy measure. First, confirmatory factor analyses at each timepoint supported a single latent factor structure (Table S1). Because participants were from two different undergraduate research programs, we evaluated the group invariance across programs at each measurement wave. The measures of self-efficacy were invariant across programs at the scalar level for T1 and T2, and partially invariant at the scalar level for T3 (the item-1 intercept exhibited non-invariance at T3; Table S2). Next, longitudinal measurement invariance was assessed at the configural, metric, and scalar levels (Widaman et al., 2010; Widaman & Reise, 1997). The results supported the scalar invariance for the self-efficacy measure across the pre/mid/post timepoints (Table S3). Thus, our longitudinal measurement and final prediction models were built with the scalar invariance constraints.

**Table S1. Confirmatory factor analysis for self-efficacy at the individual timepoints**

| Model | $\chi^2$ | *df* | *p* | RMSEA | CFI | SRMR |
|---|---|---|---|---|---|---|
| Self-efficacy$^{T1}$ | 42.766 | 8 | <.001 | .075 | .983 | .019 |
| Self-efficacy$^{T2}$ | 25.422 | 8 | <.001 | .056 | .989 | .016 |
| Self-efficacy$^{T3}$ | 24.462 | 8 | <.001 | .059 | .988 | .020 |

Notes: The residuals of self-efficacy item 1 and item 3 were allowed to correlate at T1 and T2. The residuals of self-efficacy item 3 and item 5 were allowed to correlate at T3.

**Table S2. Summary of model fit and measurement invariance test across CURE and URE groups at the individual timepoints**

| Model | $\chi^2$ | *df* | $\Delta\chi^2$ | $\Delta df$ | *p* | RMSEA | CFI | ΔCFI | SRMR | Pass? |
|---|---|---|---|---|---|---|---|---|---|---|
| **Self-efficacy$^{T1}$** | | | | | | | | | | |
| Configural | 55.566 | 16 | - | - | - | .080 | .981 | - | .021 | N/A |
| Metric | 66.257 | 21 | 9.459 | 5 | .092 | .075 | .978 | .003 | .045 | Yes |
| Scalar | 89.109 | 26 | 23.582 | 5 | <.001 | .080 | .970 | .008 | .064 | Yes |
| **Self-efficacy$^{T2}$** | | | | | | | | | | |
| Configural | 46.655 | 16 | - | - | - | .074 | .982 | - | .023 | N/A |
| Metric | 54.971 | 21 | 7.758 | 5 | .170 | .068 | .980 | .002 | .054 | Yes |
| Scalar | 71.703 | 26 | 17.182 | 5 | .004 | .071 | .973 | .007 | .067 | Yes |
| **Self-efficacy$^{T3}$** | | | | | | | | | | |
| Configural | 30.060 | 16 | - | - | - | .054 | .990 | - | .023 | N/A |
| Metric | 44.852 | 21 | 15.956 | 5 | .007 | .062 | .984 | .006 | .090 | Yes |
| Scalar | 75.508 | 26 | 34.906 | 5 | <.001 | .080 | .966 | .018 | .107 | No |
| Partial Scalar | 50.763 | 25 | 5.06 | 4 | .282 | .059 | .982 | .002 | .100 | Yes |

Notes: The residuals of self-efficacy item 1 and item 3 were allowed to correlate at T1 and T2. The residuals of self-efficacy item 3 and item 5 were allowed to correlate at T3.

**Table S3. Model comparison fit statistics for longitudinal invariance T1-T2-T3**

| Model | $\chi^2$ | df | $\Delta\chi^2$ | Δdf | p | RMSEA | CFI | ΔCFI | SRMR | Pass? |
|---|---|---|---|---|---|---|---|---|---|---|
| Configural | 580.245 | 129 | - | - | - | .068 | .932 | - | .042 | N/A |
| Metric | 601.152 | 139 | 13.852 | 10 | .181 | .066 | .931 | .001 | .047 | Yes |
| Scalar | 626.128 | 149 | 20.486 | 10 | .025 | .065 | .929 | .002 | .048 | Yes |

**S4. Four approaches to modeling scientific self-efficacy growth.** These figures reflect four approaches to modeling self-efficacy growth. In all models, $Y_{it}$ = observed score for indicator *i* at time *t*. State = true score variance. $e_{it}$ = measurement error for indicator *i* at time *t*. Trait variability that is consistent across timepoints is indicated with patterned shading ), true occasion variability in the latent factor that fluctuates across timepoints is indicated with gray shading ), latent state variables include pattern and color shading to denote conflated true trait and occasion variability, and $\varepsilon$ is the residual true variance in latent occasion variables (e.g., $\varepsilon_2$ residual true occasion variance at timepoint 2).

**Figure S1A. State-only model.** If scientific self-efficacy were repeatedly measured at the beginning, middle, and end of a research experience, total variability in the observed scientific self-efficacy scores at each timepoint (e.g., Time 1: $Y_{11}$-$Y_{61}$) is separated into latent state variance and error variances (i.e., Time 1: $\psi_1$ and $e_{11}$-$e_{61}$, respectively). Under this approach, stability is described by the degree to which latent states are correlated across timepoints.

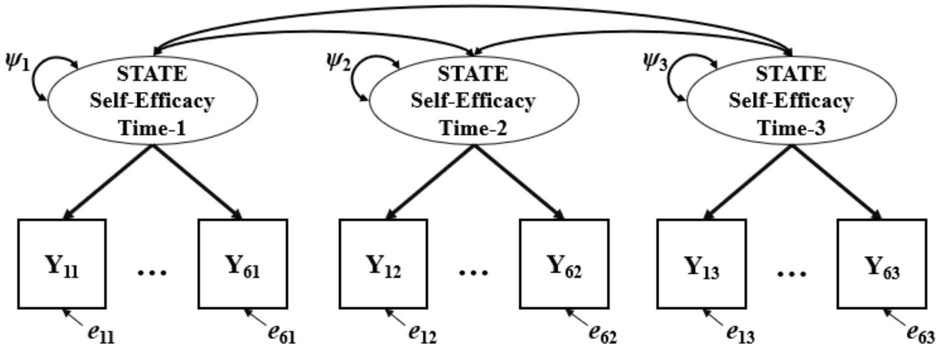


**Figure S1B. Latent state autoregressive model**. In the autoregressive model, latent state self-efficacy at a particular timepoint is predicted by prior latent state self-efficacy. The model captures stability as the influence of prior status on scientific self-efficacy (i.e., $\beta_1$ & $\beta_2$). However, recent advancements in longitudinal measurement theory show that the autoregressive approach conflates and, therefore, masks critical information about latent states and traits (Steyer et al., 2015).

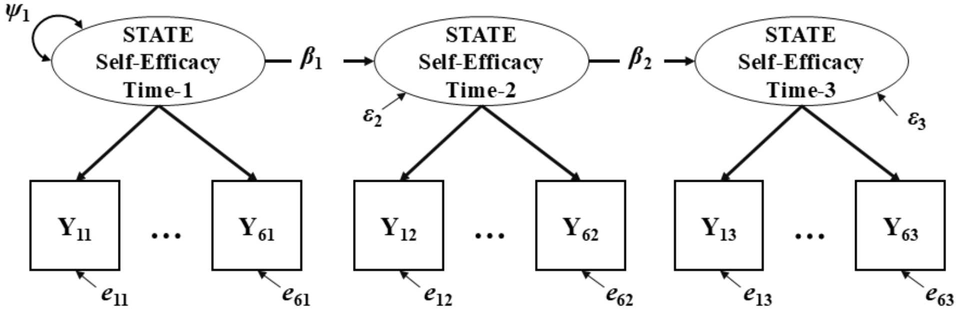

**Figures S1C and D. Trait-state-occasion models**. LST-R is a measurement theory that moves beyond the traditional distinction between "true" and "error" variances in a latent construct (Reeves & Marbach-Ad, 2016) by distinguishing between latent state, trait, and occasion aspects of "true" variance. LST-R encompasses a diverse family of longitudinal models that measure developmental processes, and LST-R subsumes the traditional confirmatory factor and autoregressive models. The trait-state-occasion (TSO) model is one of the longitudinal modeling techniques within the latent state-trait framework (Cole et al., 2005; Geiser, 2021; Steyer et al., 2015). The TSO model (Figure S1C) depicts how to capture true variability in multiple states, a single trait, and multiple occasion-specific aspects of scientific self-efficacy.

**Figure S1C. Trait-state-occasion model with one trait.**

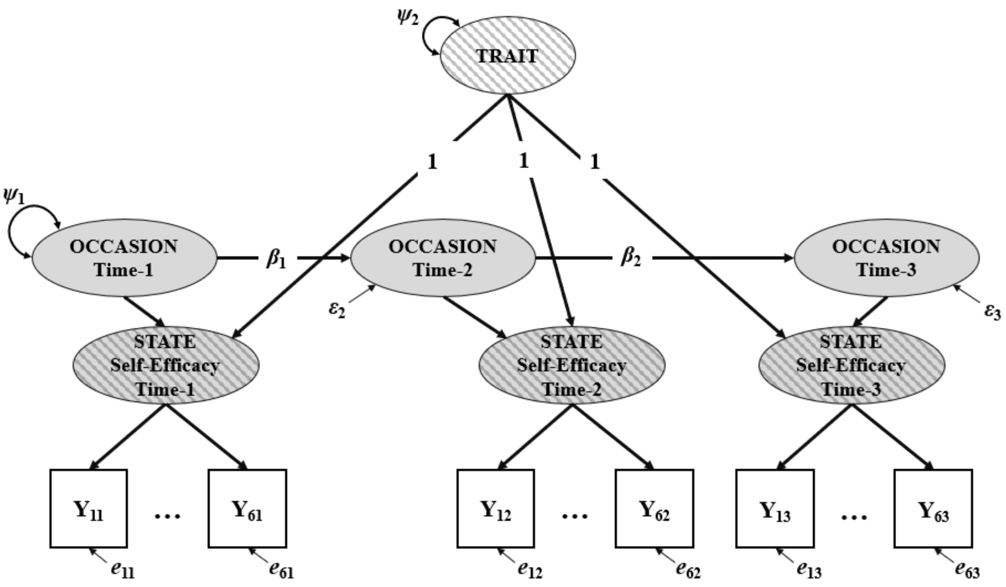


Under LST-R, a latent *state* refers to true variability in scientific self-efficacy scores measured at each timepoint, independent of measurement error variability (e.g., $e_{11}$). The total true state variability is then subdivided into portions attributable to a common trait or to specific occasions. A latent *trait* refers to the portion of true state variability in scientific self-efficacy that is stable or unchanged over time (i.e., $\psi_2$). Latent *occasions* refer to the portion of true state variability in scientific self-efficacy that fluctuates from time to time due to the situation and the person-situation interaction (e.g., $\psi_1$, $\varepsilon_2$, $\varepsilon_3$), along with possible carry-over effects from the prior occasions (i.e., $\beta_1$ & $\beta_2$).

Three important parameters are defined in the TSO model: consistency, occasion specificity, and reliability. *Consistency* refers to the proportion of variance in the observed score due to the stable trait. *Occasion specificity* refers to the proportion of the variance in the observed score due to the situation and the carry-over effects from the previous occasion. *Reliability* refers to the proportion of variance in the observed score independent of measurement error, which is the sum of the consistency and occasion specificity. These three parameters were used to inform the stability of the scientific self-efficacy measures.

When measuring a latent construct with more than one item, the method effect may exist among different items across time (e.g., due to items' unique language). The single trait TSO assumes no method effect across items. The item-specific TSO takes the method effect into account and provides less biased estimation (LaGrange & Cole, 2008; Figure S1D). In this study, the item-specific TSO model depicts how scientific self-efficacy may manifest multiple indicator-specific traits, such as a "using skills" trait distinct from a "developing theories" trait.

**Figure S1D. Trait-state-occasion model with multiple indicator-specific traits.**

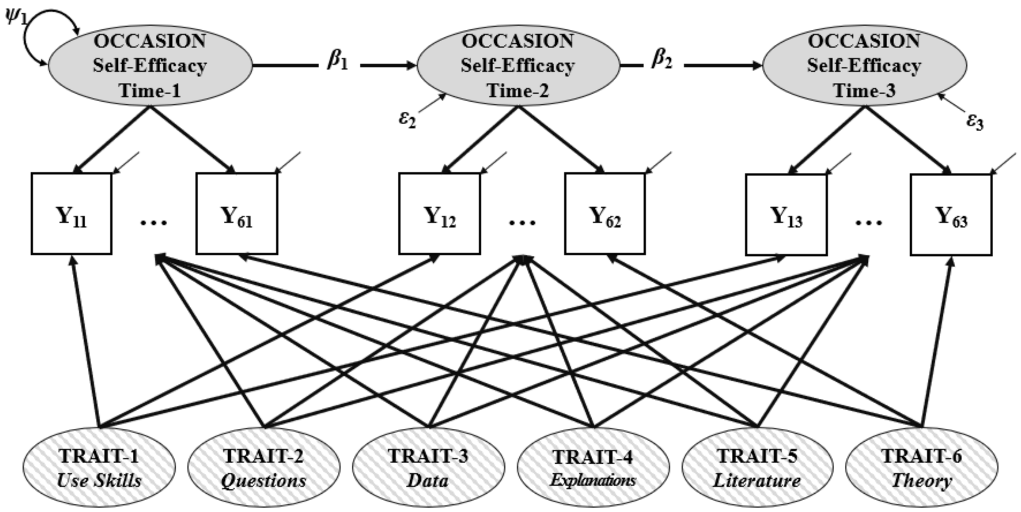

### S5. Model Comparison

All analyses were conducted in Mplus 8.10 (Muthén & Muthén, 1998-2017). We compared and evaluated six longitudinal measurement models in describing the evolvement of scientific self-efficacy over time. Models were evaluated based on the data-model fit using the following indexes and cutoff values to indicate acceptable fit: the Root-Mean-Square Error of Approximation (RMSEA $\leq .05$ or a 90% CI inclusive of .05), Comparative Fit Index (CFI $\geq .95$), and Standardized Root-Mean-Residual (SRMR $\leq .06$; Hu & Bentler, 1999). Furthermore, when comparing the alternative LST-R models, we used the Akaike Information Criterion (AIC; Akaike, 1974) and the Bayesian Information Criterion (BIC, Schwarz, 1978) to help determine which models provided the more optimal balance of fit-to-parsimony. Smaller values of AIC and BIC indicate that the models fit better in terms of parsimony. Specifically, the following six models were compared (Figure S2):

1. State model,
2. State change model,
3. Autoregressive model,
4. Trait-state-occasion model (TSO) with a single trait,
5. TSO model with multiple indicator-specific traits and equal occasion variance, and
6. TSO model with multiple indicator-specific traits and unequal occasion variance.

The assumption of whether the occasion variances are equal or not at T2 and T3 is tested in models 5 and 6 empirically. The indicator-specific trait TSO model with free estimated occasion variance (named IS-TSO model onwards) showed a significant improvement in model fit compared to other models (Table S4). To this end, the final prediction model was built on the IS-TSO model.

The final prediction model examined the relationships between predictors and the occasion components for self-efficacy at T2 and T3. Different types of tasks were negatively correlated with each other due to the constraint of a fixed total amount of research time (see Supplemental Table S5). This pattern suggested possible suppression effects (Conger, 1974; MacKinnon et al., 2000; Tzelgov & Henik, 1991). Suppression occurs when the prediction of one variable on the outcome is enhanced by the inclusion of another predictor, which leads to erroneous and misleading results. To avoid potential suppression, four different types of research tasks were included separately in the final prediction models.

**Figure S2. Six models compared.**

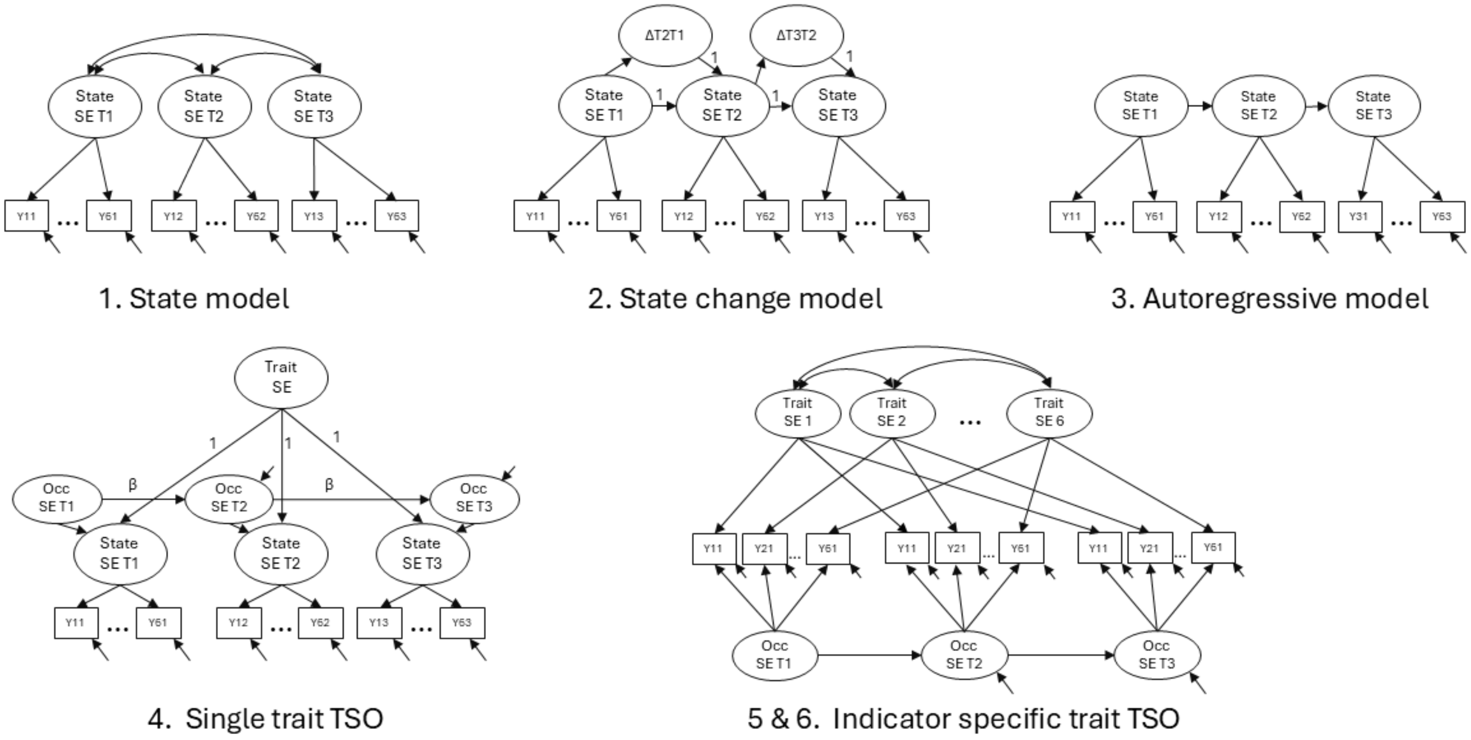

**Table S4. Measurement model comparison**

| | Model | Chi-square | df | *p* | RMSEA | CFI | SRMR | AIC | BIC |
|---|---|---|---|---|---|---|---|---|---|
| 1 | State model with invariant slopes and intercepts | 626.128 | 149 | <.001 | .065 | .929 | .048 | 30532.957 | 30718.552 |
| 2 | State change model | 626.868 | 150 | <.001 | .064 | .929 | .049 | 30532.130 | 30713.085 |
| 3 | Autoregressive model | 626.868 | 150 | <.001 | .064 | .929 | .049 | 30532.130 | 30713.085 |
| 4 | Single trait TSO with equal AR coef. & free Occ | 839.532 | 148 | <.001 | .078 | .896 | .122 | 30725.668 | 30915.903 |
| 5 | Indicator-specific trait TSO with equal Occ | 270.298 | 125 | <.001 | .039 | .978 | .080 | 30189.359 | 30486.311 |
| **6** | **Indicator-specific trait TSO with free Occ** | **252.984** | **124** | <.001 | **.037** | **.981** | **.054** | **30168.943** | **30470.131** |

Notes: AR coef. = Autoregressive coefficient; Occ = occasion variance. The bolded row indicates the best-fitting model.

### S6. Results

We first examined the bivariate correlations among the following variables: research experience type (CURE or URE), scientific self-efficacy, average number of research hours per day, standard deviation in the number of research hours per day, and involvement in one of four types of research tasks (i.e., Experimentation, Analysis, Communication, Instruction). The results show that, on average, students' research hours during the first half of the research experience (between timepoints 1 and 2, noted as T1-T2) are positively associated with students' scientific self-efficacy at the midpoint of their research experience (T2). In addition, their research hours during the second half of their research experience (midpoint to end, noted as T2-T3) show a marginal positive correlation with their scientific self-efficacy at the end of their research experience (T3). The research hour variation during T1-T2 is positively correlated with students' scientific self-efficacy at both T2 and T3. Student engagement in Analysis tasks during T1-T2 showed a positive correlation with their scientific self-efficacy at T2. Student engagement in Communication tasks during T1-T2 showed a negative correlation with their scientific self-efficacy at T2. In addition, high scientific self-efficacy at T2 was correlated with a high involvement in Instruction tasks and low involvement in Experimentation tasks later on (during T2-T3). Last, higher involvement in Experimentation during T2-T3 was correlated with lower scientific self-efficacy at T3. Table S5 shows the complete correlation matrix between variables.

The TSO model results revealed that true variability in state scientific self-efficacy development was nearly equally partitioned between multiple stable trait variances and time-specific occasion variance (Table S6). Furthermore, students' self-efficacy over time is best modeled at the indicator level – not as a single latent trait – meaning students responded differently to different items, showing distinct patterns over time regarding their specific self-efficacy beliefs. In addition, the carry-over effects between occasions are reflected in the autoregressive coefficients between adjacent occasions in the model. However, these coefficients are not significant, indicating no detectable carry-over in self-efficacy across the halves of the semester. Finally, the formal analysis was conducted to investigate the relationships between research hours, different types of research tasks, and scientific self-efficacy (Table S7).

**Table S5. Correlation matrix of outcomes and predictors at each timepoint**

| | 1 | 2 | 3 | 4 | 5 | 6 | 7 | 8 | 9 | 10 | 11 | 12 | 13 | 14 | 15 |
|---|---|---|---|---|---|---|---|---|---|---|---|---|---|---|---|
| 1. Experience | / | | | | | | | | | | | | | | |
| 2. Self-efficacy$^{T1}$ | .07* | / | | | | | | | | | | | | | |
| 3. Self-efficacy$^{T2}$ | .04 | .55*** | / | | | | | | | | | | | | |
| 4. Self-efficacy$^{T3}$ | .05 | .40*** | .67*** | / | | | | | | | | | | | |
| 5. Research Hours$^{T2}$ | .25*** | .05 | .08* | .08 | / | | | | | | | | | | |
| 6. Research Hours$^{T3}$ | .24*** | .05 | .06 | .08 | .88*** | / | | | | | | | | | |
| 7. Research Hour SD$^{T2}$ | .21*** | .05 | .13** | .13** | .67*** | .58*** | / | | | | | | | | |
| 8. Research Hour SD$^{T3}$ | .18*** | .01 | .04 | .08* | .64*** | .74*** | .72*** | / | | | | | | | |
| 9. % Analysis$^{T2}$ | .04 | .02 | .08* | .07 | .01 | .01 | .03 | -.01 | / | | | | | | |
| 10. % Instruction$^{T2}$ | -.03 | .06 | .04 | .05 | -.02 | -.04 | -.11** | -.11** | -.12** | / | | | | | |
| 11. % Communication$^{T2}$ | -.20*** | -.02 | -.11** | -.13** | -.18*** | -.13*** | -.28*** | -.18*** | -.32*** | -.20*** | / | | | | |
| 12. % Experimentation$^{T2}$ | .16*** | -.03 | .003 | .02 | .16*** | .13*** | .28*** | .23*** | -.44*** | -.40*** | -.46*** | / | | | |
| 13. % Analysis$^{T3}$ | .03 | .03 | .04 | .11** | .04 | .02 | .04 | .01 | .49*** | -.07 | -.11** | -.24*** | / | | |
| 14. % Instruction$^{T3}$ | -.26*** | .06 | .11** | .07 | -.01 | -.02 | -.08* | -.07 | .05 | .45*** | -.04 | -.30*** | -.21*** | / | |
| 15. % Communication$^{T3}$ | .04 | .02 | -.03 | -.04 | -.14*** | -.12** | -.24*** | -.20*** | -.10** | .03 | .50*** | -.36*** | -.25*** | -.22*** | / |
| 16. % Experimentation$^{T3}$ | .17*** | -.09* | -.10** | -.11** | .08* | .09* | .21*** | .20*** | -.32*** | -.35*** | -.25*** | .68*** | -.37*** | -.54*** | -.37*** |

Notes: Experience: 1 = CURE, 2 = URE. Research Hour SD = variation in research hours. %Analysis = proportion of frequency in analysis activities. %Classroom = proportion of frequency in classroom activities. %Communication = proportion of frequency in communication activities. %Experiment = proportion of frequency in experiment activities. * *p<.05,* ** *p<.01,* *** *p<.001*

**Table S6. Consistency, occasion specificity, and reliability estimates for indicators (items) across time**

| **Indicator** | 1. Use technical science skills | 2. Generate a research question to answer | 3. Figure out what data/observations to collect and how to collect them | 4. Creating explanations for results of a study | 5. Use scientific literature and/or reports to guide research | 6. Develop theories (integrate and coordinate results from multiple studies). |
|---|---|---|---|---|---|---|
| Consistency$^{T1}$ | .36 | .31 | .25 | .25 | .31 | .32 |
| Consistency$^{T2}$ | .46 | .40 | .31 | .31 | .39 | .41 |
| Consistency$^{T3}$ | .47 | .45 | .34 | .35 | .42 | .41 |
| **Mean Consistency** | **.43** | **.39** | **.30** | **.30** | **.37** | **.38** |
| **% Trait/true score** | **71%** | **53%** | **42%** | **41%** | **53%** | **52%** |
| Occasion Specificity $^{T1}$ | .21 | .39 | .50 | .51 | .39 | .43 |
| Occasion Specificity $^{T2}$ | .16 | .31 | .37 | .37 | .30 | .33 |
| Occasion Specificity $^{T3}$ | .16 | .33 | .39 | .41 | .30 | .31 |
| **Mean Specificity** | **.18** | **.34** | **.42** | **.43** | **.33** | **.36** |
| **% Occasion/true score** | **29%** | **47%** | **58%** | **59%** | **47%** | **48%** |
| Reliability $^{T1}$ | .57 | .70 | .76 | .75 | .70 | .74 |
| Reliability $^{T2}$ | .62 | .71 | .68 | .68 | .69 | .74 |
| Reliability $^{T3}$ | .63 | .78 | .73 | .76 | .72 | .72 |
| **Mean Reliability** | **.61** | **.73** | **.72** | **.73** | **.70** | **.74** |

Notes: Bolded rows indicate averages over timepoints.

**Table S7. Summary of estimates from the final model**

| Outcome | Predictor | b | SE | z | *p* | CI | β |
|---|---|---|---|---|---|---|---|
| **Model 1: Research hours and research hour variation as predictors** | | | | | | | |
| $Occ^{T2}$ | Research $hour^{T2}$ | .01 | .03 | .20 | .85 | [-.04, .05] | .01 |
| | Research hour $SD^{T2}$ | .09 | .05 | 1.66 | .10 | [-.02, .19] | .13 |
| $Occ^{T3}$ | Research $hour^{T3}$ | .01 | .02 | .37 | .71 | [-.04, .06] | .02 |
| | Research hour $SD^{T3}$ | .08 | .05 | 1.79 | .07 | [-.01, .18] | .13 |
| AR coefficient | $O_1 \rightarrow O_2$ | .20 | .13 | 1.54 | .12 | [-.05, .45] | .27 |
| AR coefficient | $O_2 \rightarrow O_3$ | .20 | .13 | 1.54 | .12 | [-.05, .45] | .19 |
| **Model 2: %Analysis as a predictor** | | | | | | | |
| $Occ^{T2}$ | Research $hour^{T2}$ | .01 | .03 | .21 | .84 | [-.04, .05] | .01 |
| | Research hour $SD^{T2}$ | .08 | .05 | 1.61 | .11 | [-.02, .18] | .12 |
| | **%Analysis $^{T2}$** | **.19** | **.09** | **2.08** | **.04** | **[.01, .38]** | **.12** |
| $Occ^{T3}$ | Research $hour^{T3}$ | .01 | .02 | .35 | .72 | [-.04, .05] | .02 |
| | Research hour $SD^{T3}$ | .08 | .05 | 1.80 | .07 | [-.01, .17] | .13 |
| | **%Analysis $^{T3}$** | **.22** | **.08** | **2.71** | **.01** | **[.06, .38]** | **.13** |
| AR coefficient | $O_1 \rightarrow O_2$ | .22 | .13 | 1.63 | .10 | [-.04. .48] | .29 |
| AR coefficient | $O_2 \rightarrow O_3$ | .22 | .13 | 1.63 | .10 | [-.04. .48] | .21 |
| **Model 3: %Communication as a predictor** | | | | | | | |
| $Occ^{T2}$ | Research $hour^{T2}$ | .01 | .03 | .34 | .73 | [-.04, .06] | .02 |
| | Research hour $SD^{T2}$ | .07 | .05 | 1.33 | .18 | [-.03, .18] | .10 |
| | %Comm $^{T2}$ | -.16 | .11 | -1.46 | .14 | [-.37, .05] | -.10 |
| $Occ^{T3}$ | Research $hour^{T3}$ | .01 | .03 | .45 | .66 | [-.04, .06] | .03 |
| | Research hour $SD^{T3}$ | .08 | .05 | 1.58 | .11 | [-.02, .17] | .12 |
| | %Comm $^{T3}$ | -.06 | .11 | -.52 | .61 | [-.26, .15] | -.03 |
| AR coefficient | $O_1 \rightarrow O_2$ | .25 | .20 | 1.29 | .20 | [-.13, .64] | .33 |
| AR coefficient | $O_2 \rightarrow O_3$ | .25 | .20 | 1.29 | .20 | [-.13, .64] | .25 |
| **Model 4: %Classroom as a predictor** | | | | | | | |
| $Occ^{T2}$ | Research $hour^{T2}$ | .01 | .03 | .22 | .83 | [-.04, .06] | .02 |
| | Research hour $SD^{T2}$ | .08 | .05 | 1.61 | .11 | [-.02, .19] | .13 |
| | %Class $^{T2}$ | -.01 | .11 | -.06 | .95 | [-.23, .22] | -.003 |
| $Occ^{T3}$ | Research $hour^{T3}$ | .01 | .02 | .36 | .72 | [-.04, .06] | .02 |
| | Research hour $SD^{T3}$ | .09 | .05 | 1.83 | .07 | [-.01, .18] | .14 |
| | %Class $^{T3}$ | .05 | .08 | .57 | .57 | [-.11, .20] | .03 |
| AR coefficient | $O_1 \rightarrow O_2$ | .20 | .13 | 1.53 | .13 | [-.06, .45] | .27 |
| AR coefficient | $O_2 \rightarrow O_3$ | .20 | .13 | 1.53 | .13 | [-.06, .45] | .19 |
| **Model 5: %Experiment as a predictor** | | | | | | | |
| $Occ^{T2}$ | Research $hour^{T2}$ | .01 | .03 | .18 | .86 | [-.05, .05] | .01 |
| | Research hour $SD^{T2}$ | .09 | .05 | 1.69 | .09 | [-.01, .19] | .13 |
| | %Experiment $^{T2}$ | -.03 | .08 | -.41 | .69 | [-.19, .12] | -.02 |
| $Occ^{T3}$ | Research $hour^{T3}$ | -.002 | .03 | -.07 | .95 | [-.05, .05] | -.004 |
| | **Research hour $SD^{T3}$** | **.11** | **.05** | **2.20** | **.03** | **[.01, .20]** | **.17** |
| | %Experiment $^{T3}$ | -.14 | .08 | -1.82 | .07 | [-.30, .01] | -.11 |
| AR coefficient | $O_1 \rightarrow O_2$ | .22 | .16 | 1.44 | .15 | [-0.08, .53] | .30 |
| AR coefficient | $O_2 \rightarrow O_3$ | .22 | .16 | 1.44 | .15 | [-0.08, .53] | .22 |

**Table S7 Notes:** Bolded rows indicate statistical significance at the conventional $p < .05$ level. Research hour = average research hour; Research hour SD = variation in research hours. %Analysis = proportion of frequency in analysis activities. %Classroom = proportion of frequency in classroom activities. %Communication = proportion of frequency in communication activities. %Experiment = proportion of frequency in experiment activities.
